# SARS-CoV-2 bearing a mutation at the S1/S2 cleavage site exhibits attenuated virulence and confers protective immunity

**DOI:** 10.1101/2021.04.29.442060

**Authors:** Michihito Sasaki, Shinsuke Toba, Yukari Itakura, Herman M. Chambaro, Mai Kishimoto, Koshiro Tabata, Kittiya Intaruck, Kentaro Uemura, Takao Sanaki, Akihiko Sato, William W. Hall, Yasuko Orba, Hirofumi Sawa

## Abstract

Severe Acute Respiratory Syndrome-Coronavirus-2 (SARS-CoV-2) possesses a discriminative polybasic cleavage motif in its spike protein that is recognized by host furin protease. Proteolytic cleavage activates the spike protein and influences both the cellular entry pathway and cell tropism of SARS-CoV-2. Here, we investigated the impact of the furin cleavage site on viral growth and pathogensis using a hamster animal model infected with SARS-CoV-2 variants bearing mutations at the furin cleavage site (S gene mutants). In the airway tissues of hamsters, the S gene mutants exhibited a low growth property. In contrast to parental pathogenic SARS-CoV-2, hamsters infected with the S gene mutants showed no body weight loss and only a mild inflammatory response, indicating the attenuated variant nature of S gene mutants. We reproduced the attenuated growth of S gene mutants in primary differenciated human airway epithelial cells. This transient infection was enough to induce protective neutralizing antibodies crossreacting with different SARS-CoV-2 lineages. Consequently, hamsters inoculated with S gene mutants showed resistance to subsequent infection with both the parental strain and the currently emerging SARS-CoV-2 variants belonging to lineages B.1.1.7 and P.1. Together, our findings revealed that the loss of the furin cleavage site causes attenuation in the airway tissues of SARS-CoV-2 and highlights the potential benefits of S gene mutants as potential immunogens.

## Introduction

Severe Acute Respiratory Syndrome-Coronavirus-2 (SARS-CoV-2) causes an infectious respiratory disease in humans, COVID-19. Patients with severe COVID-19 pneumonia exhibit high expression levels of pro-inflammatory cytokines, the so-called cytokine storm, leading to hyper-inflammation with tissue damage. Particularly, interleukin 6 (IL-6) plays a pivotal role in the hyper-inflammatory response during the acute phase of viral infection and is associated with the disease severity [1, 2]. During the global spread of SARS-CoV-2, variants carrying adaptive mutations in their spike gene have been identified in different countries, raising global concerns about disease severity, transmissibility, and escape from immunity against the ancestral SARS-CoV-2 [3-5].

Syrian hamsters and non-human primates are highly susceptible to the infection of SARS-CoV-2 and develop pneumonia with profound inflammatory responses [6-10]. Transgenic mice expressing human-ACE2 and mouse transduced human-ACE2 have also been used to investigate SARS-CoV-2 infection; however, due to the inaccessibility of mouse ACE2 [11-13], laboratory mice are resistant to infection with clinical SARS-CoV-2 strains. These animals recover from the transient infection, acquiring protective neutralizing antibodies [10, 11]. Currently, hamsters are widely used as an animal model to study pathogenicity, host immune responses, and the development of vaccines and antiviral drugs [14, 15].

The spike (S) protein of SARS-CoV-2 is a homotrimeric glycoprotein located on the virion surface and this plays a major role in virus entry into target cells by binding to specific entry receptors [16]. The S protein possesses a discriminative polybasic cleavage motif at the S1/S2 boundary which is recognized by the host furin protease and required for S protein cleavage into S1 and S2 subunits [13, 17, 18]. Importantly, this proteolytic cleavage influences the viral entry pathway (direct fusion or endocytosis) and cell tropism [17, 18]. However, our previous findings and those from other research groups suggest that SARS-CoV-2 variants bearing mutations at the furin cleavage site can be selected following passaging in Vero cells [18-25]. Although these mutants were well characterized by cell-based assays, the role of the furin cleavage site in cell tropism and pathogenicity *in vivo* remains to be elucidated. Notably, the loss of the furin cleavage site results in the attenuation of pathogenicity of SARS-CoV-2 in hamsters and human-ACE2 transgenic mice [13, 19, 25].

Here we characterized *in vivo* growth and pathogenicity of SARS-CoV-2 S gene mutants bearing deletions or substitutions at the furin cleavage sites of their S proteins [18], using a hamster model. We examined the attenuation and mild inflammatory response following infection with the S gene mutants by histopathology and cytokine expression analysis. Hamsters infected with the attenuated mutants developed neutralizing antibodies crossreacting with different lineages of SARS-CoV-2; we therefore examined whether the primary infection with an S gene mutant could protect hamster recipients from both reinfection with parental pathogenic SARS-CoV-2 and currently emerging SARS-CoV-2 variants belonging to lineages B.1.1.7 and P.1.

## Results

### Low growth properties of SARS-CoV-2 S gene mutants in Syrian hamsters

Syrian hamsters experimentally infected with SARS-CoV-2 via the intranasal route generally lose body weight until 6–7 days post infection (dpi) [7-10]. To examine the susceptibility of infection by S gene mutants, we inoculated hamsters with a clinical isolate of SARS-CoV-2, WK-521 (wild-type, WT) or S gene mutants (del2 and R685H) (Fig. 1A) [18]. Hamsters infected with WT virus showed body weight loss at 2–6 dpi, but infection with S gene mutants had no impact on the hamster body weight (Fig. 1B). The viral load of SARS-CoV-2 in hamsters reportedly decreased at 5–7 dpi [7-10]. We therefore harvested nasal turabinate and lung tissues at 4 dpi for quantification of infectious SARS-CoV-2 and its RNA. In the nasal turbinates, infectious virus titers of S gene mutants were 2–6 fold lower than those of the WT virus, whereas no difference in viral RNA levels was observed by qRT-PCR (Fig. 1C and 1D). In the lungs, the difference in growth properties between WT and S gene mutants was markedly more evident. S gene mutants produced 12–100-fold lower levels of infectious virus and viral RNA levels of S gene mutants were significantly lower than those of the WT virus (Fig. 1E and 1F). These results suggest that the S gene mutants have low pathogenicity in hamsters and have low growth capacity in hamster respiratory tissues.

**Fig 1.**
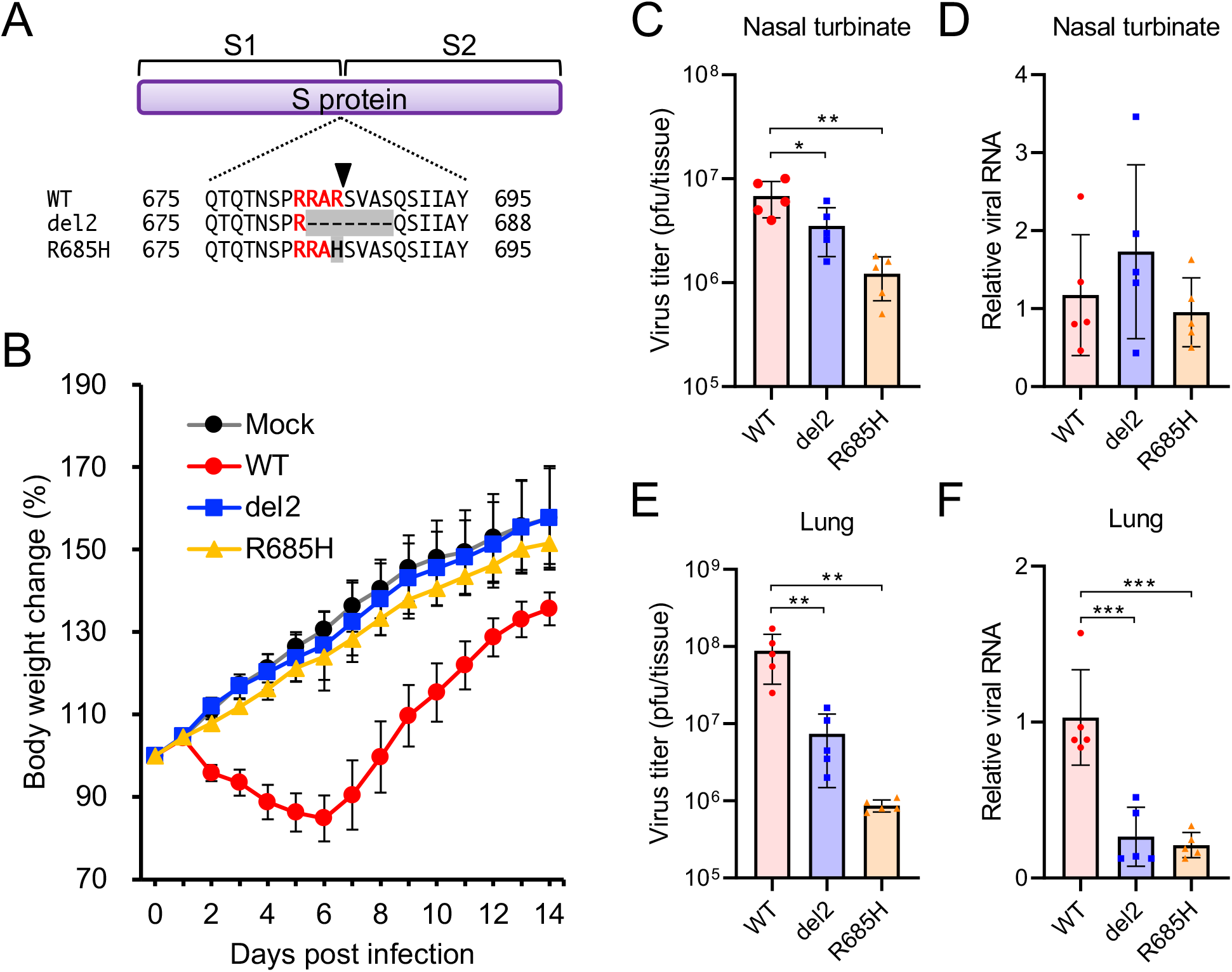
Growth of SARS-CoV-2 S gene mutants in Syrian hamsters. (A) Nascent full-length S protein is cleaved into S1 and S2 subunits at the S1/S2 cleavage site. Multiple amino acid sequence alignments were focused on the S1/S2 cleavage site of wild-type (WT) and S gene mutants (del2 and R685H). The arrowhead indicates the cleavage site. (B) Syrian hamsters were infected with SARS-CoV-2 WT or S gene mutants (del2 and R685H) via the intranasal route. Mean of body weight changes of mock- or virus-infected hamsters (*n* = 12 per group) was monitored daily. (C and E) Infectious titers in nasal turbinate (C) and lung (E) of hamsters at 4 days post infection (dpi). Viral titers in the cultures were determined using plaque assays. (D and F) Viral RNA levels relative to the WT virus in nasal turbinate (D) and lung (F) of Syrian hamsters at 4 dpi. The viral RNA levels were quantified by qRT-PCR and normalized to the expression levels of β-actin. One-way analysis of variance with Tukey’s test was used to determine the statistical significance of the differences in virus titers between the WT and S gene mutants. \**p* < 0.05, \*\**p* < 0.01, \*\*\**p* < 0.001.

### Histopathology and cytokine profiles in the lungs of hamsters infected with SARS-CoV-2 S gene mutants

We then examined gross and histological changes in the lungs of hamsters inoculated with the SARS-CoV-2 S gene mutants. On gross examination, focal pulmonary consolidations and hyperemia were observed primarily in the hilar regions of hamsters infected with WT virus at 4 dpi (Fig. 2A). In contrast, in the lungs of hamsters infected with the S gene mutants, these gross pathological changes were limited or not evident (Fig. 2A). Immunohistochemistry identified viral antigens in the nasal, bronchial and alveolar epithelia of hamsters at 2 dpi of both WT and S gene mutants (Fig. S1A and S1B). At 4 dpi, histopathological examination revealed pulmonary lesions with marked hemorrhage and inflammatory cell infiltration in the alveolar spaces of hamsters infected with the WT virus (Fig. 2B). In contrast, the histopathological changes of lungs inoculated with the S gene mutants were relatively mild compared to those of lungs inoculated with the WT virus.

**Fig 2.**
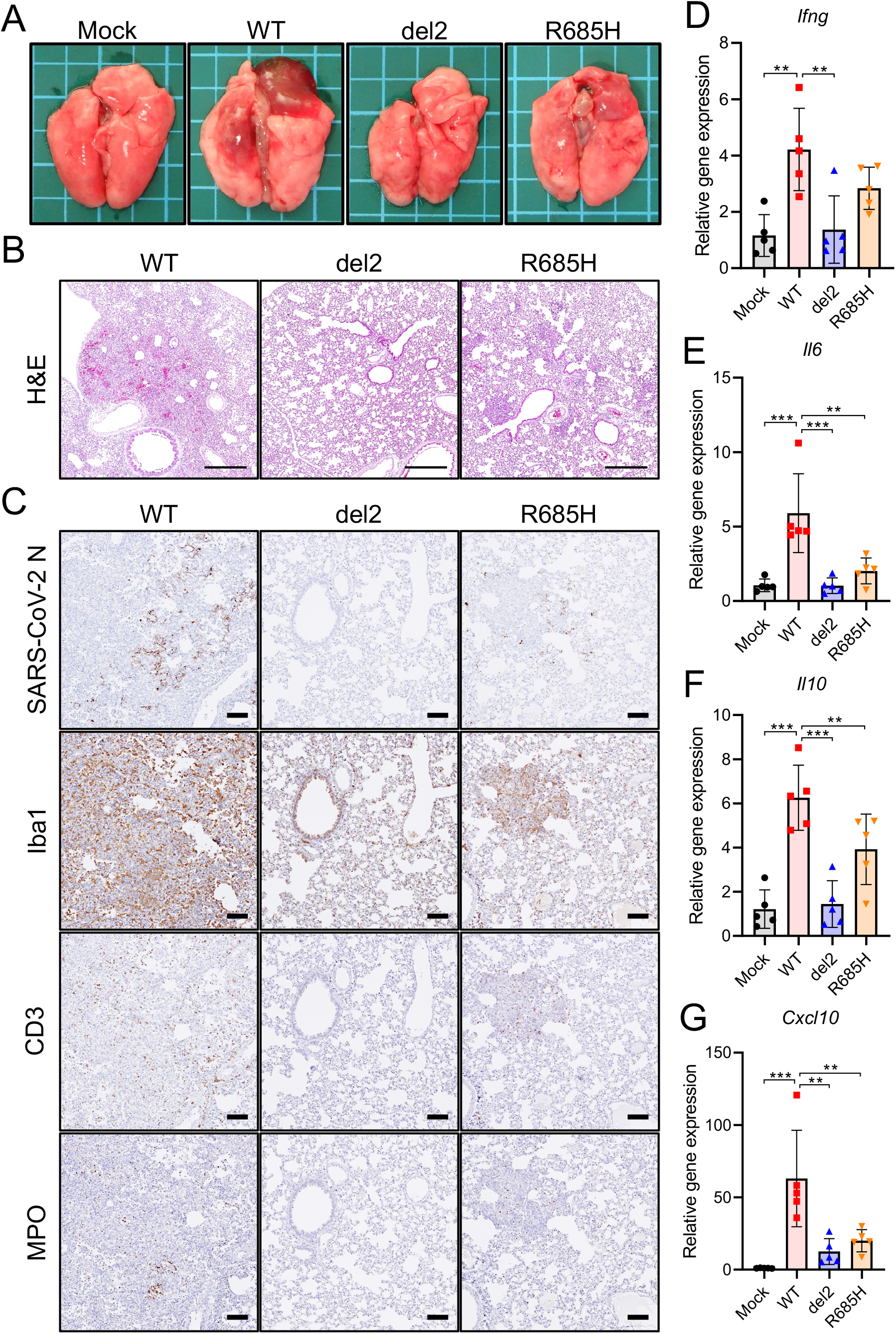
Pathological changes and immune response in lung tissues of hamsters infected with SARS-CoV-2 S gene mutants. (A) Gross pathologic images of the lungs of hamsters infected with WT or S gene mutants at 4 days post infection (dpi). (B) Histopathological images of lungs of hamsters infected with WT or S gene mutants at 4 dpi with H&E staining. Scale bars = 500 μm. (C) Immunohistochemistry for SARS-CoV-2 N protein, macrophage (Iba1), T cell (CD3) and neutrophil (MPO) markers. Cell nuclei were counterstained by hematoxylin. Scale bars = 100 μm. (D–G) Cytokine gene expression profile in lung tissues from hamsters at 4 dpi. Relative gene expression levels of indicated cytokines in the lungs as compared with lungs from mock-infected hamsters were examined by qRT-PCR. Data were normalized to β-actin. One-way analysis of variance with Tukey’s test was used to determine the statistical significance of the differences. \*\**p* < 0.01, \*\*\**p* < 0.001.

Immunohistochemistry showed widespread viral antigen-positive cells in the lung of hamsters infected with WT virus, contrasting with the relatively limited distribution of viral antigen in the lungs infected with the S gene mutants (Fig. 2C). The inflammatory cells were composed primarily of ionized calcium-binding adaptor molecule 1 (Iba1)-positive macrophages (Fig. 2C), which are thought to induce severe immune damage, consistent with observations in severe cases of COVID-19 [26, 27]. Notably, only limited inflammatory cell infiltration was observed in the lungs infected with S gene mutants (Fig. 2C).

In accordance with human COVID-19, experimental infection with SARS-CoV-2 also induced pro-inflammatory cytokine responses leading to extensive inflammatory cell infiltration in hamsters and mice [8, 12, 28]. We examined the cytokine expression levels of the hamster lungs of WT and S gene mutants at 4 dpi by qRT-PCR. WT-infection significantly upregulated the expression of IFNγ, IL-6, IL-10 and CXCL10 (also known as IP-10) in the lungs compared to that in S gene mutants (Fig. 2D–2G). These results indicate that infection with S gene mutants resulted in an attenuated inflammatory response in the lungs of the hamsters.

### Low growth property of SARS-CoV-2 S gene mutants in primary human airway epithelium

We further evaluated the growth property of S gene mutants in human airway epitheium using 3D reconstituted human nasal or bronchial epithelial cell models (nasal ECs or bronchial ECs, respectively) cultured at an air-liquid interface [29]. In control Vero E6 cells, the progeny virus titers and viral RNA levels of S gene mutants were equivalent to or higher than those of the WT virus (Fig. 3A and 3B). In contrast, the replication and growth of S gene mutants were impaired in human nasal ECs and bronchial ECs (Fig. 3C–F), consistent with the different growth properties of WT and S gene mutants in the hamster respiratory airway.

**Fig 3.**
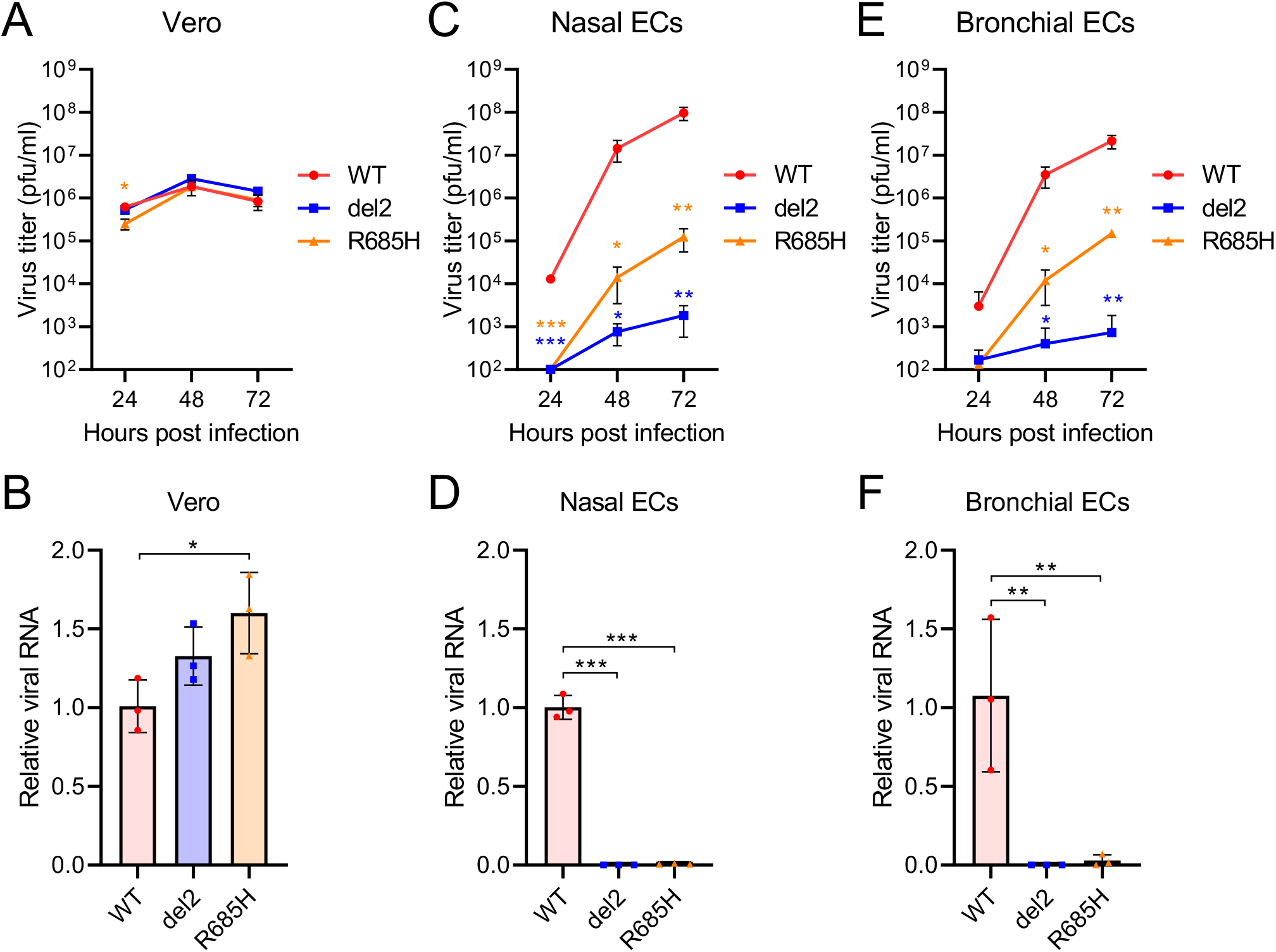
Growth of SARS-CoV-2 S gene mutants in *in vitro* cell culture. (A, C, E) Growth curves of SARS-CoV-2 WT or S gene mutants in Vero cells (A), primary human nasal epithelial cells (C), and bronchial epithelial cells (E). Viral titers in the cultures were determined using a plaque assay. (B, D, F) Viral RNA levels relative to WT virus in Vero cells (B), primary human nasal epithelial cells (D), and bronchial epithelial cells (F) at 48 h post infection. The viral RNA levels were normalized to the expression levels of β-actin. One-way analysis of variance with Tukey’s test was used to determine the statistical significance of the differences between the WT and S gene mutants. \**p* < 0.05, \*\**p* < 0.01, \*\*\**p* < 0.001.

### Infection of SARS-CoV-2 S gene mutants induces protective neutralizing antibody

Individuals infected with SARS-CoV-2 generally show detectable seroconversion at 10–14 dpi [30]. Although the virus titers in the lungs of hamsters infected with S gene mutants were lower than those with the WT virus (Fig. 1E), both hamsters infected with either WT and those infected with the S gene mutants developed similar levels of neutralizing antibody titers at 19 dpi (Fig. 4A). To investigate the protective effect of the neutralizing antibodies, we re-challenged hamsters infected with either WT or S gene mutants with the WT virus (Fig. 4B). Hamsters inoculated with WT or S gene mutants at primary infection showed no body weight loss and no macroscopic changes in the lungs following the re-challenge with the WT (WT–WT, del2-WT and R685H-WT in Fig. 4C and D). In contrast, control hamsters inoculated with PBS at the primary infection point showed marked body weight loss and macroscopic changes in the lung following secondary infection with the WT (Mock–WT in Fig. 4C and 4D). Primary infection with WT and S gene mutants prevented the proliferation of re-challenged virus and decreased viral RNA levels in nasal turbinates and lungs at 5 days post reinfection (Fig. 4E–H). In line with the inhibition of virus growth, the levels of cytokines in WT–WT, del2–WT, and R685H–WT-infected hamsters were also significantly lower than those in Mock–WT-infected hamsters (Fig. 4I–L). These results indicate that infection with the attenuated S gene mutants induced protective neutralizing antibodies and reduced disease burden during reinfection with the WT virus.

**Fig 4.**
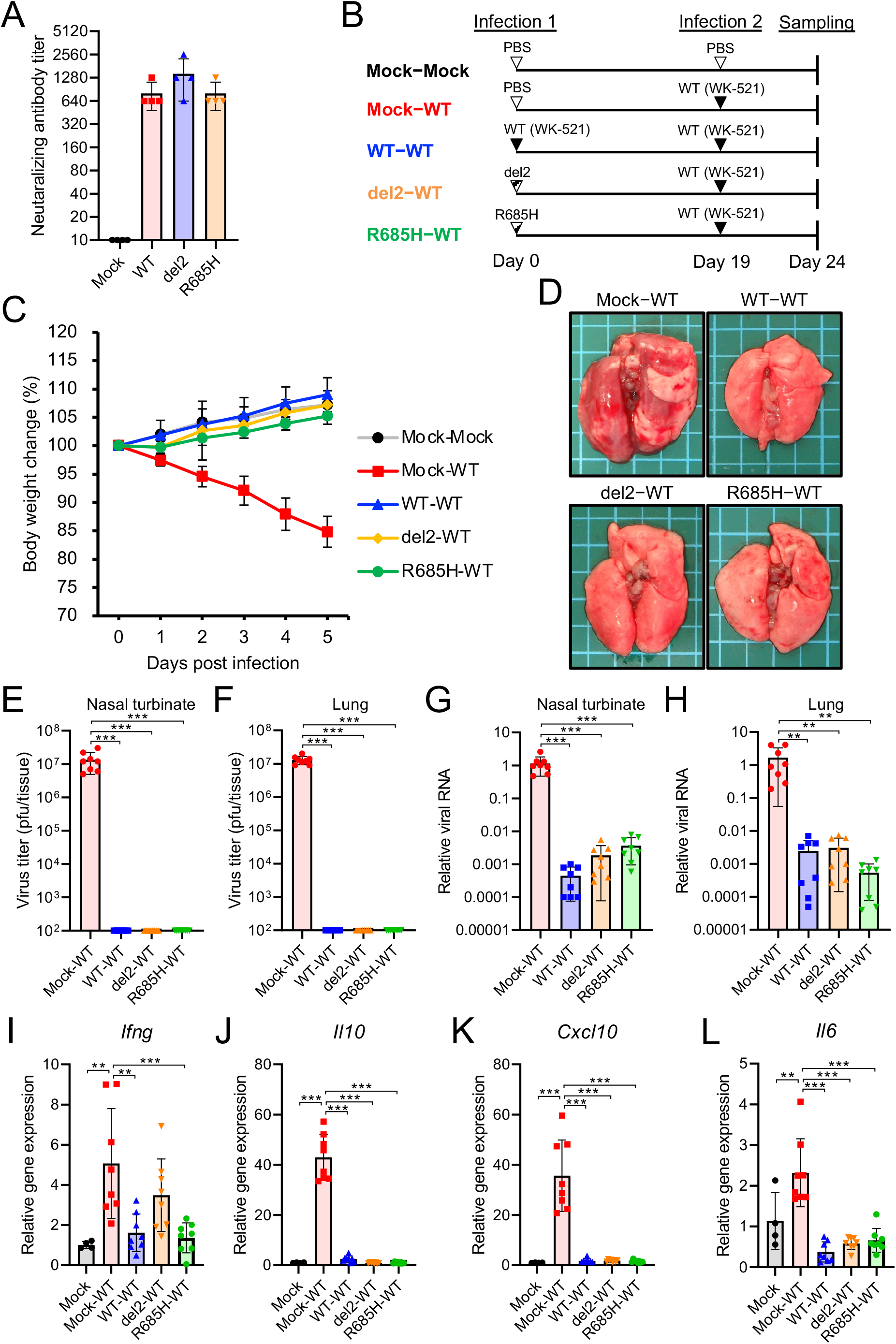
Reinfection of hamsters with SARS-CoV-2 WK-521 WT. (A) Neutalizing antibody titers in hamster serum at 19 dpi with WT or S gene mutants. (B) Schematic of primary infection, reinfection and sampling. Hamsters were infected intranasally with 1.5 × 10^4^ pfu of WT or S gene mutants. At 19 days post initial infection, hamsters were reinfected with 1.5 × 10^5^ pfu of WT virus. Mock-inoculated hamsters (Mock–Mock) and primary-infected hamsters (Mock–WT) were used as controls. (C) Mean of body weight changes of hamsters from 0 to 5 days post-reinfection. Sample sizes were *n* = 4 for the Mock–Mock group and *n* = 8 for the other groups. (D) Gross pathologic images of lungs of hamsters at 5 days post-reinfection. (E and F) Infectious virus titers in nasal turbinates (E) and lungs (F) of hamsters at 5 days post-reinfection. Viral titers in the cultures were determined using plaque assays. (G and H) Viral RNA levels relative to primary-infected hamsters (Mock–WT) in nasal turbinates (G) and lungs (H) of hamsters at 5 days post-reinfection. The viral RNA levels were quantified by qRT-PCR and normalized to the expression levels of β-actin. (I-L) Relative gene expression levels of indicated cytokines in lungs as compared with lungs from mock-infected hamsters (Mock–Mock) were examined by qRT-PCR. Data were normalized to β-actin. One-way analysis of variance with Tukey’s test was used to determine the statistical significance of the differences. \*\**p* < 0.01, \*\*\**p* < 0.001.

### Cross-reactive antibody responses to SARS-CoV-2 variants B.1.1.7 and P.1

In this study, we used S gene mutants from the SARS-CoV-2 WK-521 strain belonging to lineage A. Recently, SARS-CoV-2 variants belonging to lineages B.1.1.7 (United Kingdom), B.1.351 (South Africa), and P.1 (Brazil) have emerged. These variants possess multiple amino acid mutations in the S protein, resulting in increased transmisability and altered reactivity against neutralizing antibodies [31–35]. We utilized SARS-CoV-2 strains TY7-501 (lineage P.1) and QK002 (lineage B.1.1.7) to test whether neutralizing antibodies induced by the infection of the S gene mutant protects from infection with different SARS-CoV-2 lineages. Hamster sera in the convalescent phase of the infection of WK-521 WT or S gene mutants showed neutralizing activity against both the TY7-501 and the QK002 variants (Fig. 5A and S2A), while the cross-reactivity with TY7-501 was lower than that with QK002, presumably due to the K417T, E484K and N501Y substitutions in the S protein of TY7-501 (Fig. S3) [31–35]. We next examined whether primary infection with the WK-521 del2 mutant protects from secondary infection with TY7-501 and QK002 (Fig. 5B and S2B). Hamsters infected with del2 mutants developed no body weight loss (del2-TY7 in Fig. 5C and del2-QK002 in Fig. S2C) and no macroscopic changes in the lung at 5 days post reinfection with TY7-501 and QK002 (Fig. 5D and S2D). In the nasal turbinates and lungs of del2-TY7 and del2-QK002 hamsters, the virus titers were close to or below the detection limit of the plaque assay (Fig. 5E–F and S2E–F). The viral RNA levels were also decreased by primary infection with the del2 mutant (Fig. 5G–H and S2G–H). Consistent with the low level of virus, the expression levels of cytokines in del2-TY7 and del2-QK002 hamsters were significantly lower than those in naïve hamsters infected with SARS-CoV-2 variants (Mock-TY7 and Mock-QK002) (Fig. 5I–L and S2I–L). Our results indicate that infection with the S gene mutant del2 elicits cross-reactive immune responses to SARS-CoV-2 variants belonging to distinct lineages.

**Fig 5.**
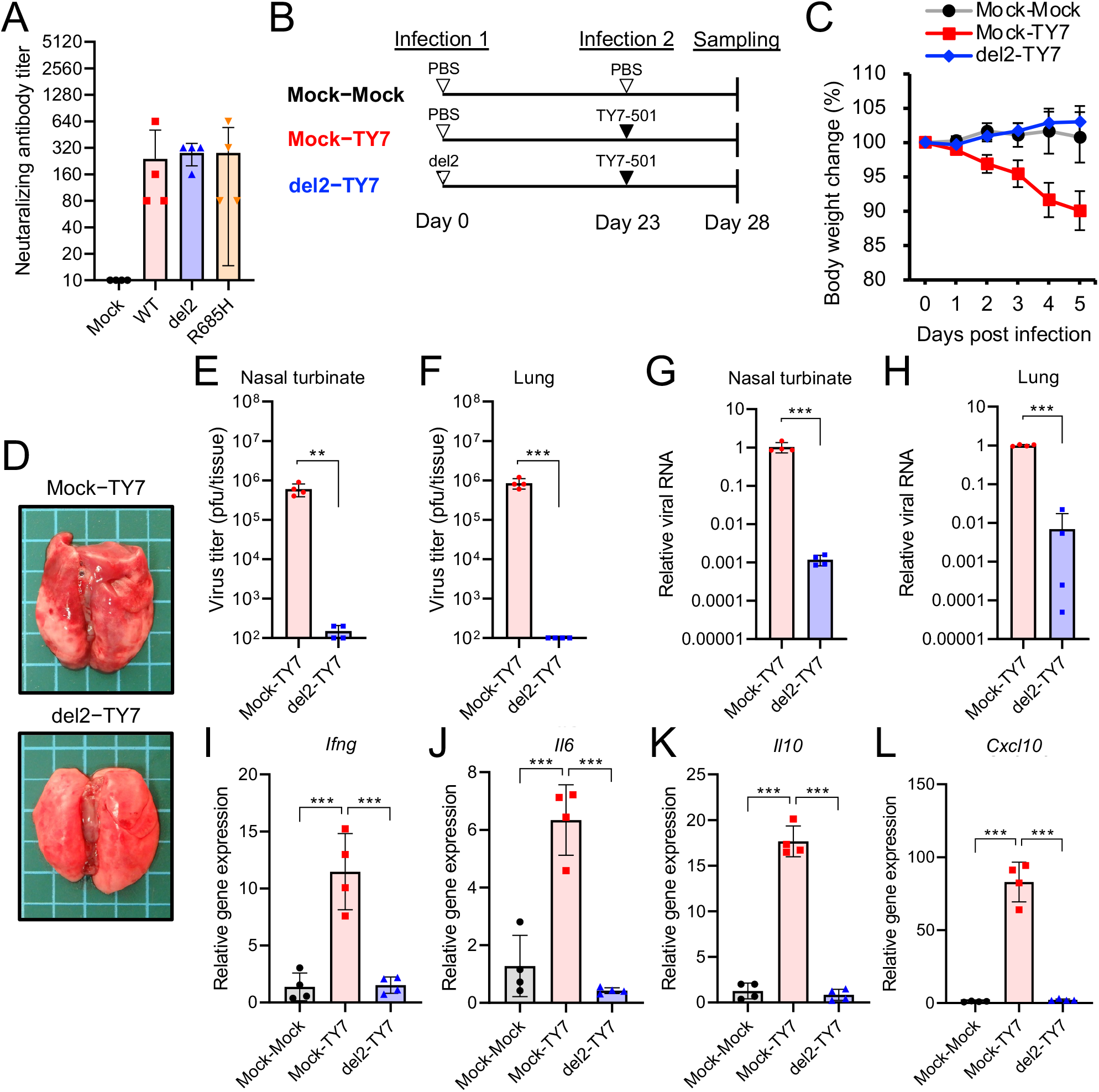
Cross-reactive neutralization among SARS-CoV-2 lineage A and lineage P.1. in hamsters. (A) Cross-neutalization test using SARS-CoV-2 TY7-501 variant (lineage P1) and hamster sera at 19 days post infection (dpi) with WT or S gene mutants of SARS-CoV-2 WK-521 (lineage A). (B) Schematic of primary infection, reinfection and sampling. Hamsters were inoculated intranasally with 1.5×10^4^ pfu of WK-521 del2 mutant or PBS. At 23 days post primary infection, hamsters were infected with 1.5 × 10^5^ pfu of TY7-501 variant. Mock-infected hamsters (Mock–Mock) and primary-infected hamsters (Mock-TY7) were used as controls. (C) Mean of body weight changes of hamsters from 0 to 5 days post-reinfection. Sample sizes were *n* = 4 for all groups. (D) Gross pathologic images of lungs of hamsters at 5 days post-reinfection. (E and F) Infectious virus titers in nasal turbinates (E) and lungs (F) of hamsters at 5 days post-reinfection. Viral titers in the cultures were determined using plaque assays. (G and H) Viral RNA levels relative to primary-infected hamsters (Mock-TY7) in nasal turbinates (G) and lungs (H) of hamsters at 5 days post-reinfection. The viral RNA levels were quantified by qRT-PCR and normalized to the expression levels of β-actin. (I-L) Relative gene expression levels of indicated cytokines in the lungs as compared with lungs from mock-infected hamsters (Mock–Mock) were examined by qRT-PCR. Data were normalized to β-actin. One-way analysis of variance with Tukey’s test was used to determine the statistical significance of the differences. \*\**p* < 0.01, \*\*\**p* < 0.001.

## Discussion

Hamsters are vulnerable to infection with SARS-CoV-2, developing pneumonia and marked body weight loss. In this study, we experimentally infected hamsters with SARS-CoV-2 clinical strains belonging to different lineages. In contrast to clinical strains, tissue culture-adapted S gene mutants bearing mutations at the S1/S2 cleavage site had limited growth capacity in hamsters, with no body weight loss and only slight lung damage, as evidenced by histopathology findings and cytokine gene expression levels. These results indicate the attenuated virulence of S gene mutants in hamsters. Other studies also reported that the loss of the furin cleavage motif at the S gene results in attenuation and ablated viral growth in hamsters and human ACE2-transgenic mice compared to the original strain [13, 19, 25]. Given the data from these studies, the low viral growth rate and subsequent mild inflammatory response in the lung tissue are characteristic hallmarks of attenuated SARS-CoV-2 variants bearing mutations at the furin cleavage site.

The cellular entry mode of S gene mutants would account for the low growth capacity of S gene mutants in the hamster airway. In the entry phase of SARS-CoV-2 infection, the S protein is primed by host TMPRSS2 or cathepsin and facilitates membrane fusion. We observed TMPRSS2 expression in the respiratory airway and impacts on the tropism of SARS-CoV-2 [36-38]. We have reported that cellular entry of S gene mutants is triggered by the cathepsin-dependent endosome pathway but not the TMPRSS2-mediated direct viral fusion at the plasma membrane [18]. The direct fusion pathway enables SARS-CoV-2 to achieve rapid cellular entry and escape from the innate immune restriction by IFN-induced transmembrane proteins (IFITMs) [39, 40]. S gene mutants thus exhibit low infectivity in certain cell lines, including human lung-derived Calu-3 cells that permit SARS-CoV-2 entry exclusively through the direct fusion pathway [17, 18]. The inability of the S gene mutants to utilize TMPRSS2 for S protein activation presumably hampers efficient virus infection and dissemination in airway epithelial cells. Nevertheless, this study demonstrated that attenuated infection is sufficient to induce a protective immunity against SARS-CoV-2 infection in hamsters.

Some attenuated virus strains—including the yellow fever virus 17D strain, measles virus Edmonston strain, poliovirus Sabin strain and varicella zoster virus Oka strain—induce protective immunity in human recipients, and have therefore been used as live-attenuated vaccines [41]. We have now demonstrated that the SARS-CoV-2 S gene mutants are attenuated variants and can induce protective immunity in hamsters. Primary infection with S gene mutants inhibited the growth of the virus in both nasal turbinates and lungs of hamsters reinfected with a pathogenic clinical strain of SARS-CoV-2. Since prophylactic administration of neutralizing IgG failed to inhibit growth of SARS-CoV-2 in nasal turbinates, this finding highlights the benefit of vaccination [42]. Moreover, inoculation with the S gene mutant del2 induced protective immunity which cross-reacted with currently emerging SARS-CoV-2 variants belonging to the lineages B.1.1.7 and P.1., which, as a result of K417T, E484K and/or N501Y mutations at RBD in the S protein, escape neutralization by some monoclonal antibodies [31-35]. This broad neutralizing activity across different lineages indicates the potential of S gene mutants as immunogens in live-attenuated vaccine candidates, although the pathogenicity of S gene mutants in humans remains to be elucidated.

Nevertheless, the recombinant SARS-CoV-2 mutant lacking the furin cleavage motif (ΔPRRA) showed low pathogenicity in human ACE2-transgenic mice as well as hamsters [13]. In humans, naturally arising SARS-CoV-2 variants lacking the furin cleavage motif have been identified as minor populations of quasispecies in clinical specimens from COVID-19 patients [43]. In primary differenciated human epithelial cells, we have demonstrated the low growth properties of S gene mutants. These observations suggest that SARS-CoV-2 mutants lacking the furin cleavage site can infect human airway, albeit with low growth properties. Further *in vivo* studies using non-human primates will provides more insights on the implications and pathogenicity of S gene mutants. In conclusion, the findings in the present study show the potential of developing live-attenuated vaccines for the prevention of SARS-CoV-2 infection.

## Methods

### Cells

Vero (Vero E6, ATCC, Manassas, VA) and Vero-TMPRSS2 [18] cells were maintained in Dulbecco’s Modified Eagle’s Medium (DMEM) supplemented with 10% fetal bovine serum (FBS). Differentiated human nasal and bronchial epithelial cells (nasal ECs and bronchial ECs) in an ALI-culture were obtained as MucilAir-nasal and MucilAir-bronchial, and maintained in MucilAir culture medium (all from Epithelix, Genève, Switzerland). All cells were incubated at 37°C with 5% CO_2_.

### Viruses

SARS-CoV-2 WK-521 (EPI_ISL_408667), QK002 (EPI_ISL_768526), TY7-501 (EPI_ISL_833366) strains were provided by Drs. Saijyo, Shimojima and Ito (National Institute of Infectious Diseases, Japan); the original stock of these virus strains was prepared by inoculation of Vero-TMPRSS2 cells. S gene mutant virus clones of WK-521, del2 and R685H, were isolated as previously described and propagated in Vero cells [18].

### Ethical statement

All of the animal experiments were performed in accordance with the National University Corporation, Hokkaido University Regulations on Animal Experimentation. The protocol was reviewed and approved by the Institutional Animal Care and Use Committee of Hokkaido University (approval no. 20-0060).

### Hamster infection

For virological and histopathological analyses in single infection, 4-to 6-week-old male Syrian hamsters (Japan SLC, Shizuoka, Japan) were inoculated intranasally with 1.5×10^4^ plaque forming units (pfu) of wild-type WK-521 (WT), del2 or R685H viruses in 200 μl of PBS. Body weights of the infected hamsters were monitored daily. At 2, 4 or 19 dpi, a subset of the infected hamsters were euthanized under deep anesthesia by isoflurane inhalation, and tissue samples (nasal turbinate, lung and blood) were harvested.

For the reinfection experiments, 4-week-old male Syrian hamsters were inoculated intranasally with 1.5×10^4^ pfu of WT, del2 or R685H viruses in 200 μl of PBS or PBS only (mock-infected controls). At 19 or 23 dpi, the hamsters were reinfected with 1.5 × 10^5^ pfu of WT, QK002 or TY7-501 virus strains in 200 μl of PBS. At 5 days post reinfection (24 or 28 days post initial infection), tissue samples (nasal turbinate and lung) were harvested.

Whole lung and nasal turbinate tissues were homogenized in PBS with TissueRuptor (Qiagen, Hilden, Germany). A part of the homogenate was centrifuged for 2 min at 2,310 × *g* to pellet tissue debris and the supernatant was subjected to plaque assays using Vero-TMPRSS2 cells for virus titration as previously described [18]. The remaining part of the homogenate was mixed with TRIzol LS (Invitrogen; Thermo Fisher Scientific, Waltham, MA) and subjected to RNA extraction with Direct-zol RNA MiniPrep kit (Zymo Research, Irvine, CA). For relative quantification of viral RNA and host mRNAs, cDNA was synthesized with SuperScript IV VILO Master Mix (Invitrogen) and analyzed by qRT-PCR with Probe qPCR Mix (Takara, Kusatsu, Japan). Target RNA levels were normalized to hamster β-actin and and calculated by the ΔΔCt method. Primers and probes were previsouly described and are listed in Table S1 [44, 45].

### Histopathology and immunohistochemistry

Nasal turbinate and lung tissue samples were harvested from hamsters infected at 2 or 4 dpi of SARS-CoV-2. Tissue samples were fixed in 10% phosphate-buffered formalin and nasal turbinate were decalcified with 10% EDTA solution (pH 7.0). Tissue samples were then embedded in paraffin. The paraffin blocks were sectioned at 4 μm thickness and mounted on Platinum PRO micro glass slides (Matsunami, Osaka, Japan). For histopathological analysis, slides were stained with hematoxylin and eosin (H&E). For immunohistochemical analysis, slides were heated in citrate buffer for 5 min using a pressure cooker for antigen retrieval and blocking with Block Ace (KAC, Kyoto, Japan), followed by staining with anti-SARS-CoV-2 spike antibody (GTX632604, GeneTex, Hsinchu, Taiwan), anti-SARS-CoV-2 nucleocapsid antibody (GTX635679, GeneTex), anti-CD3 (ab16669, Abcam, Cambridge, UK), anti-MPO (A039829-2, DAKO; Agilent Santa clara, CA) or anti-Iba1 (019-19741, FUJIFILM Wako, Osaka, Japan).

Immunostaining was detected by EnVision system peroxidase-labeled anti-rabbit or anti-mouse immunoglobulin (DAKO) and visualized with a Histofine diaminobenzidine substrate kit (Nichirei Biosciences, Tokyo, Japan).

### Infection and growth of SARS-CoV-2 in *in vitro* cell culture

Human nasal ECs and bronchial ECs in an ALI-culture were infected at the apical surface with either WT, del2 or R685H viruses at an MOI of 0.1. After 1 h of incubation, apical area of cells were washed three times with PBS and then cells were maintained under ALI-culture conditions. At 24, 48, and 72 h post infection (hpi), 200 μl of culture medium was added at the apical side and the fluid was harvested for virus titration after 20 min incubation. Vero cells were infected with either WT, del2 or R685H viruses at an MOI of 0.01. After 1 h of incubation, cells were washed three times with PBS and then cultured in fresh medium with 2% FBS. The culture supernatants were harvested at 24, 48 and 72 hpi. Virus titers were determined by plaque assays as previously described [18]. For viral RNA quantification, RNA was extracted with Direct-zol RNA MiniPrep kit at 48 hpi (Vero cells) or 72 hpi (Human nasal ECs and bronchial ECs) and analyzed by qRT-PCR with the Thunderbird Probe One-step Probe qRT-PCR Kit (Toyobo, Osaka, Japan). Viral RNA levels were normalized to non-human primate β-actin (Table S1) or human β-actin (Hs99999903_m1, Applied Biosystems; Thermo Fisher Scientific) and calculated by the ΔΔCt method [46].

### Virus neutralization assays

Serum samples were collected from hamsters at 19 dpi after infection with the SARS-CoV-2 WK-521 strain and heat-inactivated at 56°C for 30 min. Serial two-fold dilutions of serum samples in DMEM containing 2% FBS were incubated with 160 pfu of SARS-CoV-2 WK-521, QK002 or TY7-501 strains at 37°C for 1 h. The serum-virus mixtures were then add to Vero-TMPRSS2 cells in 96 well plates. After 4 dpi, viral cytopathic effects were examined under an inverted microscope. The neutralization titer was defined as the reciprocal of the highest serum dilution that completely inhibited the cytopathic effect.

### Statistical analysis

Data were expressed as the mean ±SD. Statistical analysis was performed by One-way analysis of variance (ANOVA) with Tukey’s test using GraphPad Prism 8 (GraphPad Prism Software, San Diego, CA).

### Role of the funding source

The funders had no role in study design, data collection, data analysis, data interpretation, or writing of this article.

## Acknowledgments

We thank Drs. Saijyo, Shimojima and Ito at National Institute of Infectious Diseases, Japan for providing SARS-CoV-2 WK-521, QK002, TY7-501 strains. This work was supported by the Japan Agency for Medical Research and Development (AMED) under Grant numbers JP21wm0125008, PJ21wm0225003, PJ21fk0108104, PJ20fk0108509 and PJ20fk0108509, and Scientific Research on Innovative Areas from the Ministry of Education, Culture, Sports, Science and Technology (MEXT) of Japan under Grant numbers 16H06429, 16H06431 and 16K21723, and Japan Science and Technology Agency (JST) Moonshot R&D under Grant numbers JPMJMS2025..

## Competing interests

The authors S.T., K.U., T.S., and A.S. are employees of Shionogi & Co., Ltd. Other authors declare no competing interests.

---

**Fig S1.Viral antigen-positive cells in hamsters at 2 day post infection.** Immunohistochemistry of SARS-CoV-2 N in nasal turbinates (A) and lungs (B) at 2 days post infection (dpi) of SARS-CoV-2 WK-521 WT or S gene mutants. Cell nuclei were counterstained with hematoxylin. Scale bars = 100 μm.

**Fig S2. Cross-reactive neutralization among SARS-CoV-2 lineage A and lineage B.1.1.7 in hamsters** (A) Cross-neutalization test using SARS-CoV-2 QK002 variant (lineage B.1.1.7) and hamster sera at 19 days post infection (dpi) with WT or S gene mutants of SARS-CoV-2 WK-521 (lineage A). (B) Schematic representation of primary infection, reinfection and sampling. Hamsters were inoculated intranasally with 1.5 × 10^4^ pfu of WK-521 del2 mutant or PBS. At 23 days post primary infection, hamsters were infected with 1.5 × 10^5^ pfu of QK002 variant. Mock-infected hamsters (Mock–Mock) and primary-infected hamsters (Mock-QK002) were used as controls. Mock–Mock hamsters are the same individuals as those represented in Fig.4. (C) Mean of body weight changes of hamsters from 0 to 5 days post-reinfection. Sample sizes were *n* = 4 for all groups. (D) Gross pathologic images of lungs of hamsters at 5 days post-reinfection. (E and F) Infectious virus titers in nasal turbinates (E) and lungs (F) of hamsters at 5 days post-reinfection. Viral titers in the cultures were determined using plaque assays. (G and H) Viral RNA levels relative to primary-infected hamsters (Mock-QK002) in nasal turbinates (G) and lungs (H) of hamsters at 5 days post-reinfection. The viral RNA levels were quantified by qRT-PCR and normalized to the expression levels of β-actin. (I-L) Relative gene expression levels of indicated cytokines in lungs compared with lungs from mock-infected hamsters (Mock–Mock) were examined by qRT-PCR. Data were normalized to β-actin. One-way analysis of variance with Tukey’s test was used to determine the statistical significance of the differences. \*\**p* < 0.01, \*\*\**p* < 0.001.

**Fig S3. Multiple amino acid sequence alignment of S protein of SARS-CoV-2** Multiple sequence alignment based on the full length of the deduced S protein sequence of WK521 (lineage A), QK002 (lineage B.1.1.7) and TY7-501 (lineage P.1.). Amino acid substitutions and deletions in QK002 and TY7-501 are shown as pink and green boxes, respectively.

