## Supplemental Figure S1-S3 for "SARS-CoV-2 bearing a mutation at the S1/S2 cleavage site exhibits attenuated virulence and confers protective immunity"

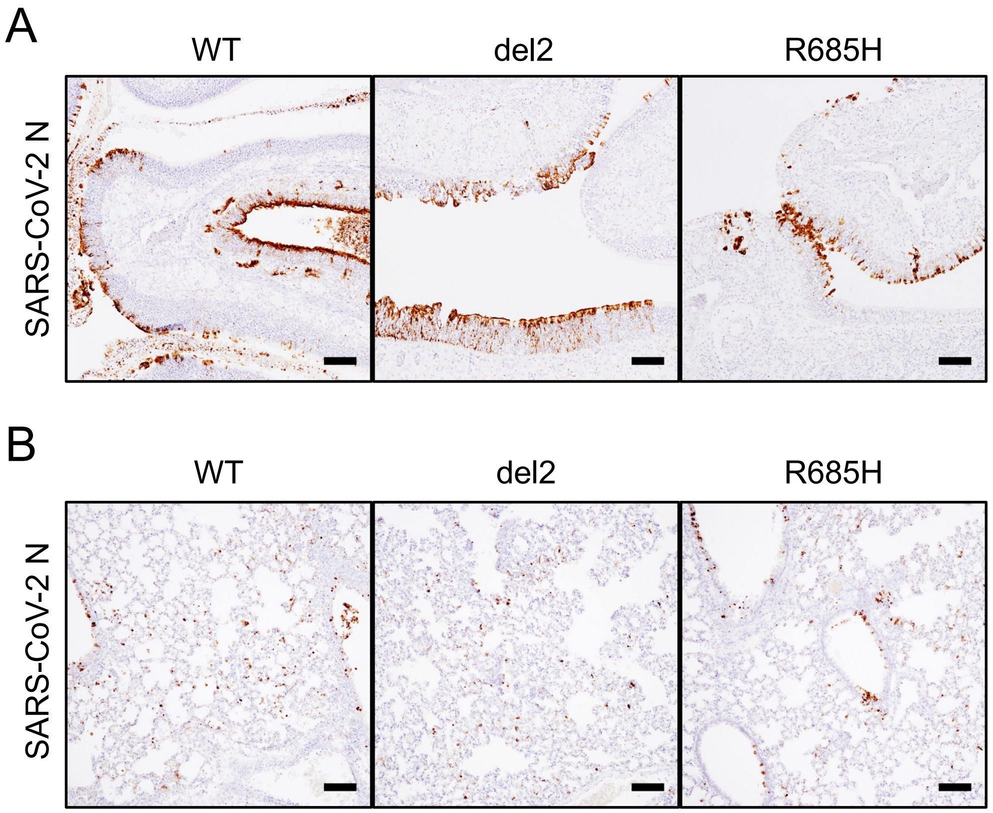

### Figure S2

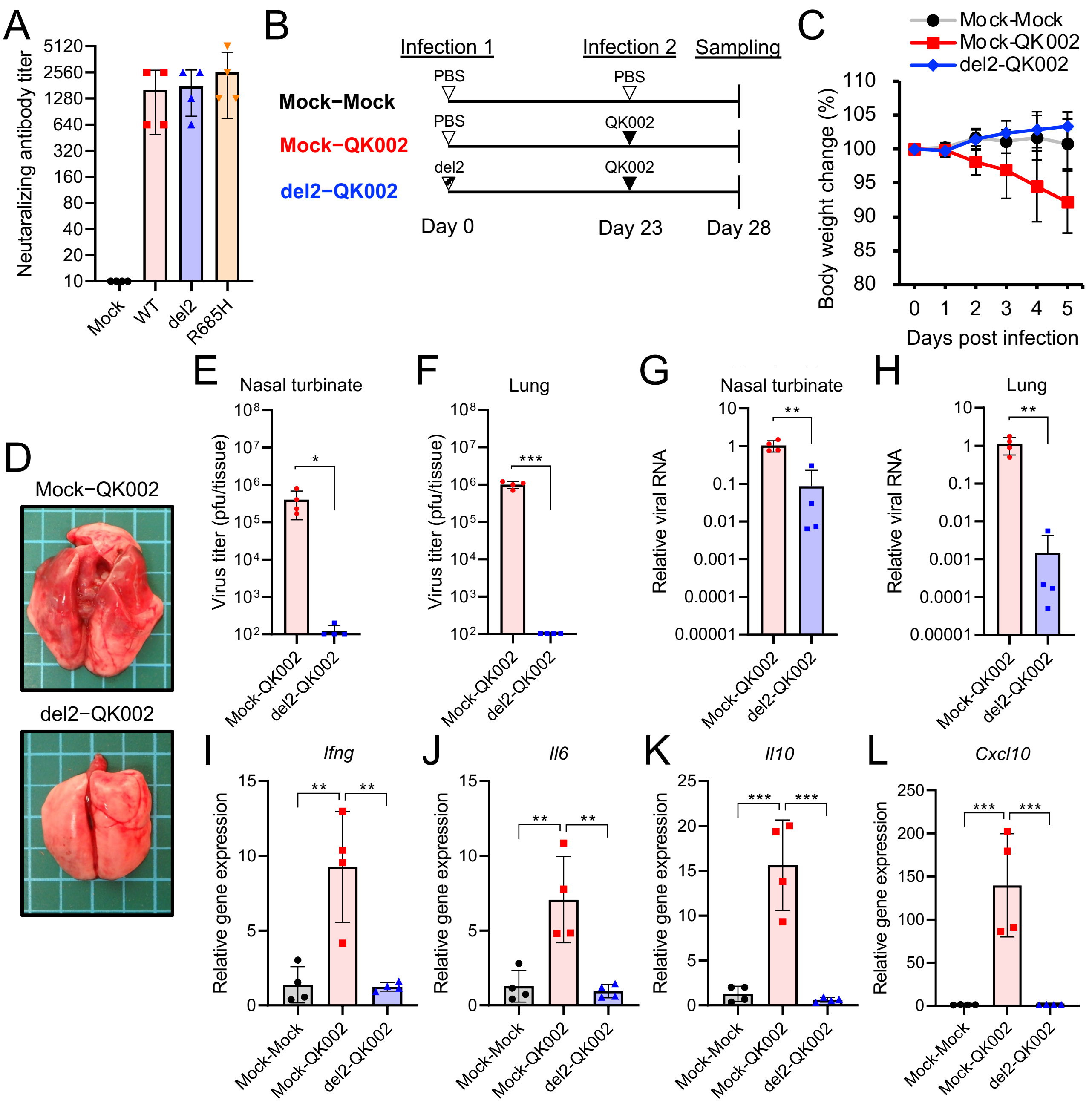

A
Neutaralizing antibody titer
5120
2560
1280
640
320
160
80
40
20
10
Mock
WT
del2
R685H
B
Infection 1
Infection 2
Sampling
Mock−Mock
Mock−QK002
del2−QK002
PBS
PBS
PBS
QK002
del2
QK002
Day 0
Day 23
Day 28
C
Mock-Mock
Mock-QK002
del2-QK002
Body weight change (%)
110
105
100
95
90
85
80
0 1 2 3 4 5
Days post infection
D
Mock−QK002
del2−QK002
E
Nasal turbinate
Virus titer (pfu/tissue)
*
Mock-QK002
del2-QK002
F
Lung
***
G
Nasal turbinate
Relative viral RNA
**
H
Lung
**
I
Ifng
Relative gene expression
Mock-Mock
Mock-QK002
del2-QK002
J
Il6
K
Il10
L
Cxcl10

### Figure S3

```
WK-521     1 MFVFLVLLPLVSSQCVNLTTRTQLPPAYTNSFTRGVYYPDKVFRSSVLHSTQDLFLPFFSNVTWFHAIHVSGTNGTKRFDNPVLPFNDGV
QK002      1 MFVFLVLLPLVSSQCVNLTTRTQLPPAYTNSFTRGVYYPDKVFRSSVLHSTQDLFLPFFSNVTWFHAI--SGTNGTKRFDNPVLPFNDGV
TY7-501    1 MFVFLVLLPLVSSQCVNFTNRTQLPSAYTNSFTRGVYYPDKVFRSSVLHSTQDLFLPFFSNVTWFHAIHVSGTNGTKRFDNPVLPFNDGV

WK-521    91 YFASTEKSNIIRGWIFGTTLDSKTQSLLIVNNATNVVIKVCEFQFCNDPFLGVYYHKNNKSWMESEFRVYSSANNCTFEYVSQPFLMDLE
QK002     89 YFASTEKSNIIRGWIFGTTLDSKTQSLLIVNNATNVVIKVCEFQFCNDPFLGVY-HKNNKSWMESEFRVYSSANNCTFEYVSQPFLMDLE
TY7-501   91 YFASTEKSNIIRGWIFGTTLDSKTQSLLIVNNATNVVIKVCEFQFCNYPFLGVYYHKNNKSWMESEFRVYSSANNCTFEYVSQPFLMDLE

WK-521   181 GKQGNFKNLREFVFKNIDGYFKIYSKHTPINLVRDLPQGFSALEPLVDLPIGINITRFQTLLALHRSYLTPGDSSSGWTAGAAYYVGYL
QK002    178 GKQGNFKNLREFVFKNIDGYFKIYSKHTPINLVRDLPQGFSALEPLVDLPIGINITRFQTLLALHRSYLTPGDSSSGWTAGAAYYVGYL
TY7-501  181 VKQGNFKNLSEFVFKNIDGYFKIYSKHTPINLVRDLPQGFSALEPLVDLPIGINITRFQTLLALHRSYLTPGDSSSGWTAGAAYYVGYL

WK-521   271 QPRTFLLKYNENGTITDAVDCALDPLSETKCTLKSFTVEKGIYQTSNFRVQPTESIVRFPNITNLCPFGEVFNATRFASVYAWNRKRISN
QK002    268 QPRTFLLKYNENGTITDAVDCALDPLSETKCTLKSFTVEKGIYQTSNFRVQPTESIVRFPNITNLCPFGEVFNATRFASVYAWNRKRISN
TY7-501  271 QPRTFLLKYNENGTITDAVDCALDPLSETKCTLKSFTVEKGIYQTSNFRVQPTESIVRFPNITNLCPFGEVFNATRFASVYAWNRKRISN

                                     Receptor binding domain (RBD)
WK-521   361 CVADYSVLYNSASFSTFKCYGVSPTKLNDLCFTNVYADSFVIRGDEVRQIAPGQTGKIADYNYKLPDDFTGCVIAWNSNNLDSKVGGNYN
QK002    358 CVADYSVLYNSASFSTFKCYGVSPTKLNDLCFTNVYADSFVIRGDEVRQIAPGQTGKIADYNYKLPDDFTGCVIAWNSNNLDSKVGGNYN
TY7-501  361 CVADYSVLYNSASFSTFKCYGVSPTKLNDLCFTNVYADSFVIRGDEVRQIAPGQTGTIADYNYKLPDDFTGCVIAWNSNNLDSKVGGNYN

                                             484                 501
WK-521   451 YLYRLFRKSNLKPFERDISTEIYQAGSTPCNGVEGFNCYFPLQSYGFQPTNGVGYQPYRVVVLSFELLHAPATVCGPKKSTNLVKNKCVN
QK002    448 YLYRLFRKSNLKPFERDISTEIYQAGSTPCNGVEGFNCYFPLQSYGFQPTYGVGYQPYRVVVLSFELLHAPATVCGPKKSTNLVKNKCVN
TY7-501  451 YLYRLFRKSNLKPFERDISTEIYQAGSTPCNGVKGFNCYFPLQSYGFQPTYGVGYQPYRVVVLSFELLHAPATVCGPKKSTNLVKNKCVN

WK-521   541 FNFNGLTGTGVLTESNKKFLPFQQFGRDIADTTDAVRDPQTLEILDITPCSFGGVSVITPGTNTSNQVAVLYQDVNCTEVPVAIHADQLT
QK002    538 FNFNGLTGTGVLTESNKKFLPFQQFGRDIDDTTDAVRDPQTLEILDITPCSFGGVSVITPGTNTSNQVAVLYQGVNCTEVPVAIHADQLT
TY7-501  541 FNFNGLTGTGVLTESNKKFLPFQQFGRDIADTTDAVRDPQTLEILDITPCSFGGVSVITPGTNTSNQVAVLYQGVNCTEVPVAIHADQLT

                                                         Furin cleavage site
WK-521   631 PTWRVYSTGSNVFQTRAGCLIGAEHVNNSYECDIPIGAGICASYQTQTNSPRRARSVASQSIIAYTMSLGAENSVAYSNNSIAIPTNFTI
QK002    628 PTWRVYSTGSNVFQTRAGCLIGAEHVNNSYECDIPIGAGICASYQTQTNSHRRARSVASQSIIAYTMSLGAENSVAYSNNSIAIPINFTI
TY7-501  631 PTWRVYSTGSNVFQTRAGCLIGAEYVNNSYECDIPIGAGICASYQTQTNSPRRARSVASQSIIAYTMSLGAENSVAYSNNSIAIPTNFTI

WK-521   721 SVTTEILPVSMTKTSVDCTMYICGDSTECSNLLLQYGSFCTQLNRALTGIAVEQDKNTQEVFAQVKQIYKTPPIKDFGGFNFSQILPDPS
QK002    718 SVTTEILPVSMTKTSVDCTMYICGDSTECSNLLLQYGSFCTQLNRALTGIAVEQDKNTQEVFAQVKQIYKTPPIKDFGGFNFSQILPDPS
TY7-501  721 SVTTEILPVSMTKTSVDCTMYICGDSTECSNLLLQYGSFCTQLNRALTGIAVEQDKNTQEVFAQVKQIYKTPPIKDFGGFNFSQILPDPS

WK-521   811 KPSKRSFIEDLLFNKVTLADAGFIKQYGDCLGDIAARDLICAQKFNGLTVLPPLLTDEMIAQYTSALLAGTITSGWTFGAGAALQIPFAM
QK002    808 KPSKRSFIEDLLFNKVTLADAGFIKQYGDCLGDIAARDLICAQKFNGLTVLPPLLTDEMIAQYTSALLAGTITSGWTFGAGAALQIPFAM
TY7-501  811 KPSKRSFIEDLLFNKVTLADAGFIKQYGDCLGDIAARDLICAQKFNGLTVLPPLLTDEMIAQYTSALLAGTITSGWTFGAGAALQIPFAM

WK-521   901 QMAYRFNGIGVTQNVLYENQKLIANQFNSAIGKIQDSLSSTASALGKLQDVVNQNAQALNTLVKQLSSNFGAISSVLNDILSRLDKVEAE
QK002    898 QMAYRFNGIGVTQNVLYENQKLIANQFNSAIGKIQDSLSSTASALGKLQDVVNQNAQALNTLVKQLSSNFGAISSVLNDILARLDKVEAE
TY7-501  901 QMAYRFNGIGVTQNVLYENQKLIANQFNSAIGKIQDSLSSTASALGKLQDVVNQNAQALNTLVKQLSSNFGAISSVLNDILSRLDKVEAE

WK-521   991 VQIDRLITGRLQSLQTYVTQQLIRAAEIRASANLAATKMSECVLGQSKRVDFCGKGYHLMSFPQSAPHGVVFLHVTYVPAQEKNFTTAPA
QK002    988 VQIDRLITGRLQSLQTYVTQQLIRAAEIRASANLAATKMSECVLGQSKRVDFCGKGYHLMSFPQSAPHGVVFLHVTYVPAQEKNFTTAPA
TY7-501  991 VQIDRLITGRLQSLQTYVTQQLIRAAEIRASANLAAIKMSECVLGQSKRVDFCGKGYHLMSFPQSAPHGVVFLHVTYVPAQEKNFTTAPA

WK-521  1081 ICHDGKAHFPREGVFVSNGTHWFVTQRNFYEPQIITTDNTFVSGNCDVVIGIVNNTVYDPLQPELDSFKEELDKYFKNHTSPDVDLGDIS
QK002   1078 ICHDGKAHFPREGVFVSNGTHWFVTQRNFYEPQIITTHNTFVSGNCDVVIGIVNNTVYDPLQPELDSFKEELDKYFKNHTSPDVDLGDIS
TY7-501 1081 ICHDGKAHFPREGVFVSNGTHWFVTQRNFYEPQIITTDNTFVSGNCDVVIGIVNNTVYDPLQPELDSFKEELDKYFKNHTSPDVDLGDIS

WK-521  1171 GINASVVNIQKEIDRLNEVAKNLNESLIDLQELGKYEQYIKWPWYIWLGFIAGLIAIVMVTIMLCCMTSCCSCLKGCCSCGSCCKFDEDD
QK002   1168 GINASVVNIQKEIDRLNEVAKNLNESLIDLQELGKYEQYIKWPWYIWLGFIAGLIAIVMVTIMLCCMTSCCSCLKGCCSCGSCCKFDEDD
TY7-501 1171 GINASFVNIQKEIDRLNEVAKNLNESLIDLQELGKYEQYIKWPWYIWLGFIAGLIAIVMVTIMLCCMTSCCSCLKGCCSCGSCCKFDEDD

WK-521  1261 SEPVLKGVKLHYT
QK002   1258 SEPVLKGVKLHYT
TY7-501 1261 SEPVLKGVKLHYT
```
