## Supplemental Table S1 for "SARS-CoV-2 bearing a mutation at the S1/S2 cleavage site exhibits attenuated virulence and confers protective immunity"

**Supplementary Material**

**Table S1. Sequence information of the primers and probes used for qRT-PCR in this study**

| Target name | Primer/Probe | Sequence (5' to 3') | Reference |
| --- | --- | --- | --- |
| SARS-CoV-2 N | Primer | 5'-AAATTTTGGGGACCAGGAAC-3' | Shirato *et al*. 2020 |
|  | Primer | 5'-TGGCAGCTGTGTAGGTCAAC-3' | [44] |
|  | Probe | 5'-FAM-ATGTCGCGCATTGGCATGGA-BHQ1-3' |  |
| β-actin (hamster) | Primer | 5'-ACTGCCGCATCCTCTTCCT-3' | Zivcec *et al*. 2011 |
|  | Primer | 5'-TCGTTGCCAATGGTGATGAC-3' | [45] |
|  | Probe | 5'-FAM-CCTGGAGAAGAGCTATGAGCTGCCTGATG-BHQ1-3' |  |
| Ifng (hamster) | Primer | 5'-GGCCATCCAGAGGAGCATAG-3' | Zivcec *et al*. 2011 |
|  | Primer | 5'-TTTCTCCATGCTGCTGTTGAA-3' | [45] |
|  | Probe | 5'-FAM-CACCATCAAGGCAGACCTGTTTGCTAACTT-BHQ1-3' |  |
| Il1b (hamster) | Primer | 5'-GGCTGATGCTCCCATTCG-3' | Zivcec *et al*. 2011 |
|  | Primer | 5'-CACGAGGCATTTCTGTTGTTCA-3' | [45] |
|  | Probe | 5'-FAM-CAGCTGCACTGCAGGCTCCGAG-BHQ1-3' |  |
| Il4 (hamster) | Primer | 5'-CCACGGAGAAAGACCTCATCTG-3' | Zivcec *et al*. 2011 |
|  | Primer | 5'-GGGTCACCTCATGTTGGAAATAAA-3' | [45] |
|  | Probe | 5'-FAM-CAGGGCTTCCCAGGTGCTTCGCAAGT-BHQ1-3' |  |
| Il6 (hamster) | Primer | 5'-CCTGAAAGCACTTGAAGAATTCC-3' | Zivcec *et al*. 2011 |
|  | Primer | 5'-GGTATGCTAAGGCACAGCACACT-3' | [45] |
|  | Probe | 5'-FAM-AGAAGTCACCATGAGGTCTACTCGGCAAAA-BHQ1-3' |  |
| Il10 (hamster) | Primer | 5'-GTTGCCAAACCTTATCAGAAATGA-3' | Zivcec *et al*. 2011 |
|  | Primer | 5'-TTCTGGCCCGTGGTTCTCT-3' | [45] |
|  | Probe | 5'-FAM-CAGTTTTACCTGGTAGAAGTGATGCCCCAGG-BHQ1-3' |  |
| Tnfa (hamster) | Primer | 5'-GGAGTGGCTGAGCCATCGT-3' | Zivcec *et al*. 2011 |
|  | Primer | 5'-AGCTGGTTGTCTTTGAGAGACATG-3' | [45] |
|  | Probe | 5'-FAM-CCAATGCCCTCCTGGCCAACG-BHQ1-3' |  |
| Cxcl10 (hamster) | Primer | 5'-GCCATTCATCCACAGTTGACA-3' | Zivcec *et al*. 2011 |
|  | Primer | 5'-CATGGTGCTGACAGTGGAGTCT-3' | [45] |
|  | Probe | 5'-FAM-CGTCCCGAGCCAGCCAACGA-BHQ1-3' |  |
| β-actin (primate) | Primer | 5'-ACCCCAAGGCCAACCG-3' | Overbergh *et al*. 2005 |
|  | Primer | 5'-ACAGCCTGGATGGCCACRTACA-3' | [46] |
|  | Probe | 5'-FAM-TGACCCAGATCATGTTT-MGB-3' | with modification |
